# *MALAT1* expression indicates cell quality in single-cell RNA sequencing data

**DOI:** 10.1101/2024.07.14.603469

**Authors:** Zoe A. Clarke, Gary D. Bader

**Affiliations:** Department of Molecular Genetics, University of Toronto, Toronto, Ontario, Canada; The Donnelly Centre, University of Toronto, Toronto, Ontario, Canada; Department of Computer Science, University of Toronto, Toronto, Ontario, Canada; Lunenfeld-Tanenbaum Research Institute, Toronto, Ontario, Canada; Princess Margaret Research Institute, University Health Network, Toronto, Ontario, Canada; CIFAR Multiscale Human Program, CIFAR, Toronto, Ontario, Canada

**Keywords:** single-cell RNA-sequencing, quality control, MALAT1, empty droplets, filtering, splicing, lncRNA

## Abstract

Single-cell RNA sequencing (scRNA-seq) has revolutionized our understanding of cell types and tissues. However, empty droplets and poor quality cells are often captured in single cell genomics experiments and need to be removed to avoid cell type interpretation errors. Many automated and manual methods exist to identify poor quality cells or empty droplets, such as minimum RNA count thresholds and comparing the gene expression profile of an individual cell to the overall background RNA expression of the experiment. A versatile approach is to use unbalanced overall RNA splice ratios of cells to identify poor quality cells or empty droplets. However, this approach is computationally intensive, requiring a detailed search through all sequence reads in the experiment to quantify spliced and unspliced reads. We found that the expression level of *MALAT1,* a non-coding RNA retained in the nucleus and ubiquitously expressed across cell types, is strongly correlated with this splice ratio measure and thus can be used to similarly identify low quality cells in scRNA-seq data. Since it is easy to visualize the expression of a single gene in single-cell maps, *MALAT1* expression is a simple cell quality measure that can be quickly used during the cell annotation process to improve the interpretation of cells in tissues of human, mouse and other species with a conserved *MALAT1* function.

## Introduction

The analysis of biological signals in single-cell RNA sequencing (scRNA-seq) data relies on the assumptions that all measured cells are intact and that the gene expression data associated with each cell comes solely from that cell. The methods used to support these assumptions depend on the protocol used for cell isolation and transcriptome sequencing. For droplet-based methods, cells may release RNA transcripts into the solution of the experiment during cell processing creating ambient RNA which makes it difficult to tell which transcripts come from which cells (Young and Behjati 2018; Heaton et al. 2020). Ambient RNA can also fill an empty droplet making it look like a real cell (Lun et al. 2019; Muskovic and Powell 2021; Montserrat-Ayuso and Esteve-Codina 2024). Certain tissues and cell types are more prone to this effect, as some cell types characteristically have a higher rate of rupturing and releasing their cellular RNA during the cell isolation process. In such experiments, droplets may contain damaged cells, cell fragments, or ambient cytoplasmic RNA, and still pass all traditional cell filtering steps (Muskovic and Powell 2021).

A droplet-based scRNA-seq experiment typically produces hundreds of thousands of droplets, a small proportion of which contain cells. Cell-containing droplets are identified if they have high UMI counts or have moderate UMI counts and are different compared to a background RNA profile learned from all droplets in the experiment. The results are plotted on a “knee” plot showing the read count distribution per droplet (Zheng et al. 2017; Lun et al. 2019). Following this, droplets that contain multiple cells (“doublets”) (McGinnis et al. 2019; Wolock et al. 2019; Bais and Kostka 2020) or stressed or damaged cells with high mitochondrial content (Ilicic et al. 2016) are often removed from the data. However, even with these filters applied, many single-cell datasets suffer from contamination challenges, with damaged cells and ambient RNA-filled empty droplets being retained and eventually called as intact cells (Muskovic and Powell 2021; Schmauch et al. 2022; Braun et al. 2023; Macnair and Robinson 2023; Montserrat-Ayuso and Esteve-Codina 2024). DropletQC was developed to remove damaged cells from droplet-based experiments (Muskovic and Powell 2021). This method identifies damaged cells if their cytoplasmic (spliced mRNA) to nuclear (intron-retaining mRNA) ratio, called the “nuclear fraction”, is unbalanced. If droplets have few intronic reads, or a low nuclear fraction, they likely don’t contain a nucleus and are considered empty and removed from the sample. Conversely, if the nuclear fraction is high, droplets are labeled as “damaged cells” with a potentially ruptured plasma membrane allowing cytoplasmic, but not nuclear, mRNA to leak out into solution. Although DropletQC is helpful in interpreting cell quality in scRNA-seq data, it requires a computationally-intensive search of all DNA sequencing reads aligned to the genome (contained in BAM files) to identify intronic reads. In some public data sets, it is challenging to get access to the raw sequence data needed to compute the nuclear fraction to support reanalysis. Since the creation of DropletQC, additional methods have taken advantage of a cell’s splice ratio to improve cell filtering. For example, SampleQC recognizes the typical Gaussian distribution of features within cell types (including the splice ratio), and flags outliers (Macnair and Robinson 2023), and QClus similarly analyzes multiple quality metrics in single-nucleus RNA-seq data and flags cells with few unspliced reads (Schmauch et al. 2022).

*MALAT1* is a highly abundant long non-coding RNA (lncRNA) that is retained in the nucleus (Sun and Ma 2019). It is consistently highly expressed across mammalian species, and its function is related to splicing regulation, the cell cycle, or the structure of the nuclear speckle (Hutchinson et al. 2007; Sun and Ma 2019). The DropletQC authors used *MALAT1*’s expression level to assess whether a droplet contains a nucleus (Muskovic and Powell 2021) and others have shown that *MALAT1*’s expression level correlates with the nuclear fraction (Montserrat-Ayuso and Esteve-Codina 2024). We analyzed a variety of scRNA-seq data to identify genes that correlate with nuclear fraction and found that *MALAT1* expression is the most generally correlated and that a low *MALAT1* expression level can be used to flag these for further investigation and likely filtering. We conclude that *MALAT1* expression is a useful technical signal that easily and quickly indicates the presence of a nucleus in droplets in scRNA-seq data and recommend that this measure be incorporated into scRNA-seq analysis pipelines.

## Results

### MALAT1 expression is an indicator of cell quality

We compared the nuclear fractions of healthy and diseased datasets from human and mouse with normalized *MALAT1* expression and found these values are highly correlated (Table 1, Figure 1). *MALAT1* is the top correlated gene in 81% of samples, and is always highly correlated (Table 1). We also isolated empty droplets not included in the Cell Ranger filtered cell matrix in all 10x Genomics-provided datasets listed in Table 1 and performed differential gene expression between the empty droplets and cells as called by Cell Ranger (Zheng et al. 2017). *MALAT1* is always the top differentially expressed gene in the cells (p-val < 1.73 e-84 for all six 10x Genomics samples tested where we used filtered and raw matrices directly output by Cell Ranger).

**Figure 1.**
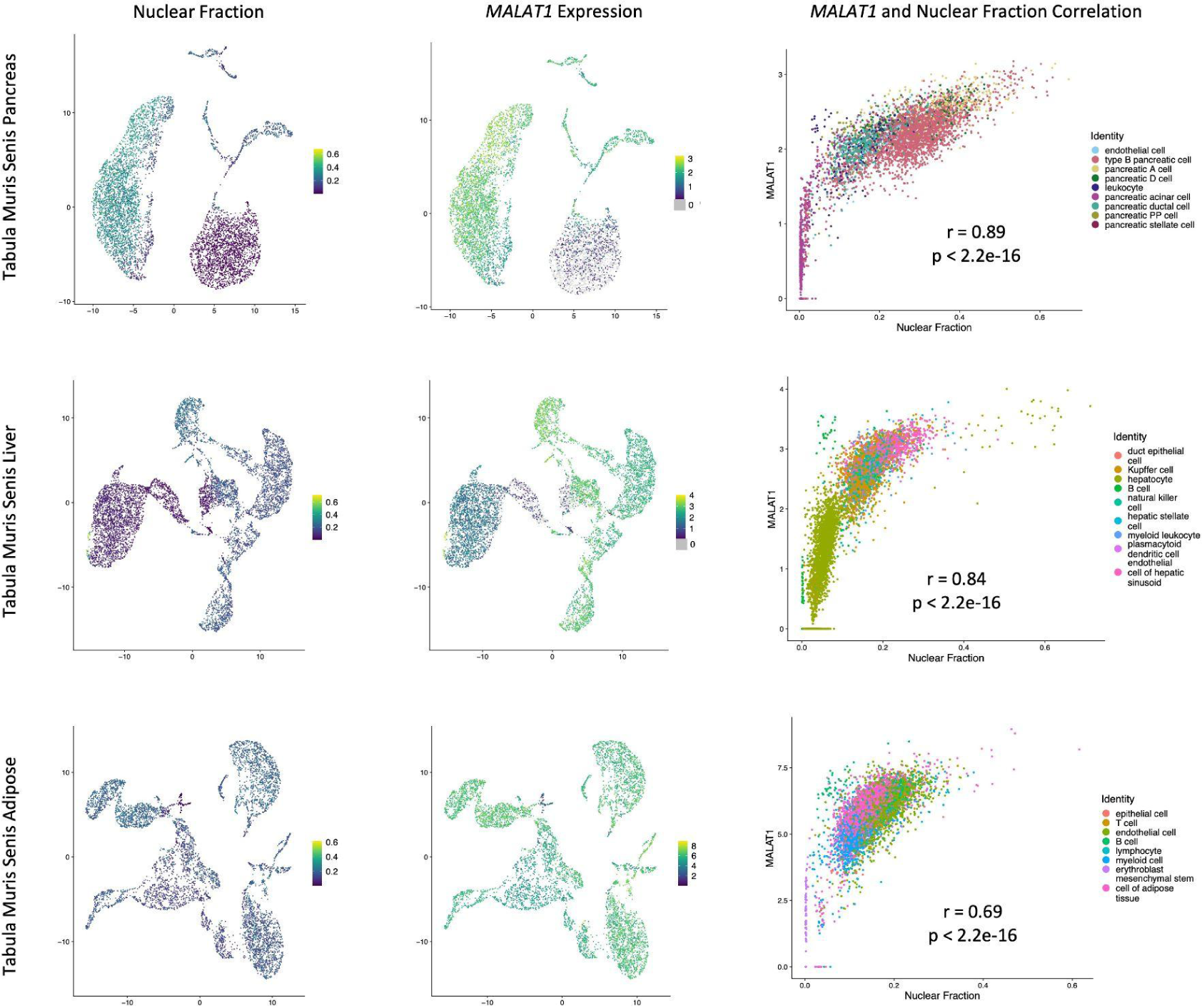
Nuclear fraction values are strongly correlated with normalized *MALAT1* expression values. In the first column, nuclear fraction values are visualized as colours on the UMAPs of various datasets (described in Table 1). Lighter colours represent higher nuclear fraction values, and darker colours represent lower values. The second column contains normalized *MALAT1* expression visualized as colours on UMAP coordinates, with the same colour scheme. The third column contains scatterplots of nuclear fraction values and normalized *MALAT1* expression values, with the correlation coefficients and p-values indicated by r and p respectively. Cells are coloured by cell type as identified by the publishers of the dataset.

**Table 1:**
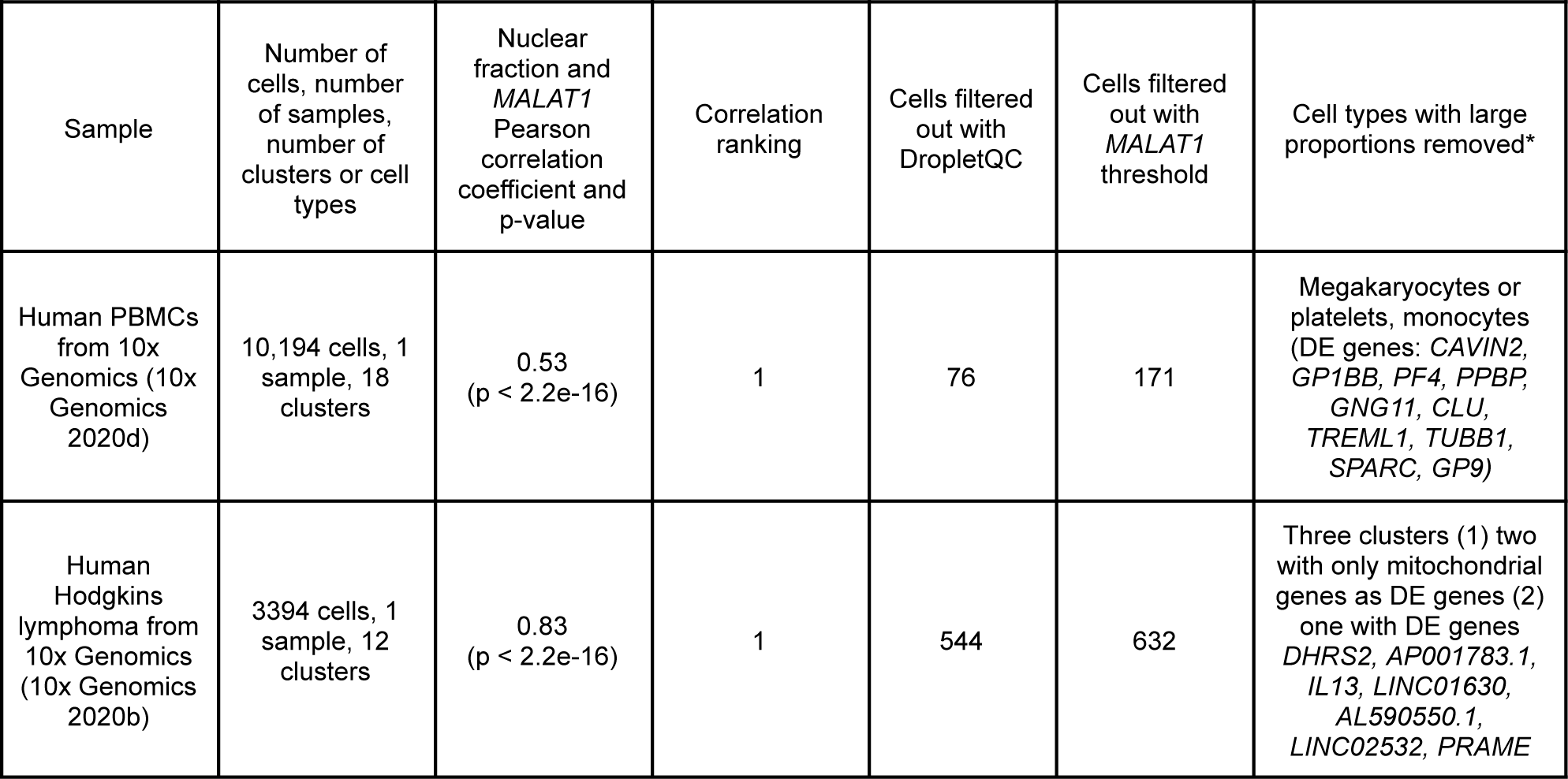
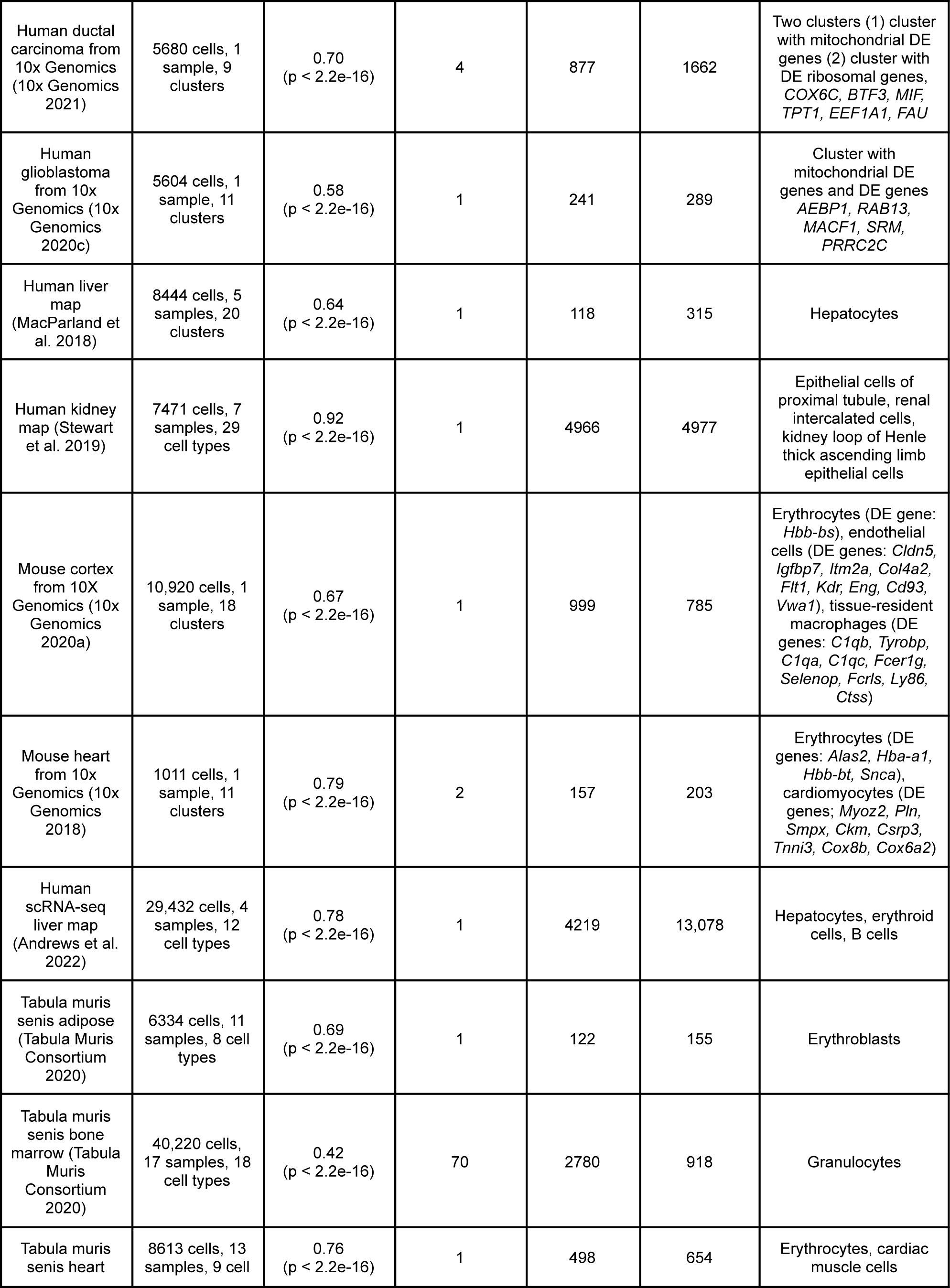
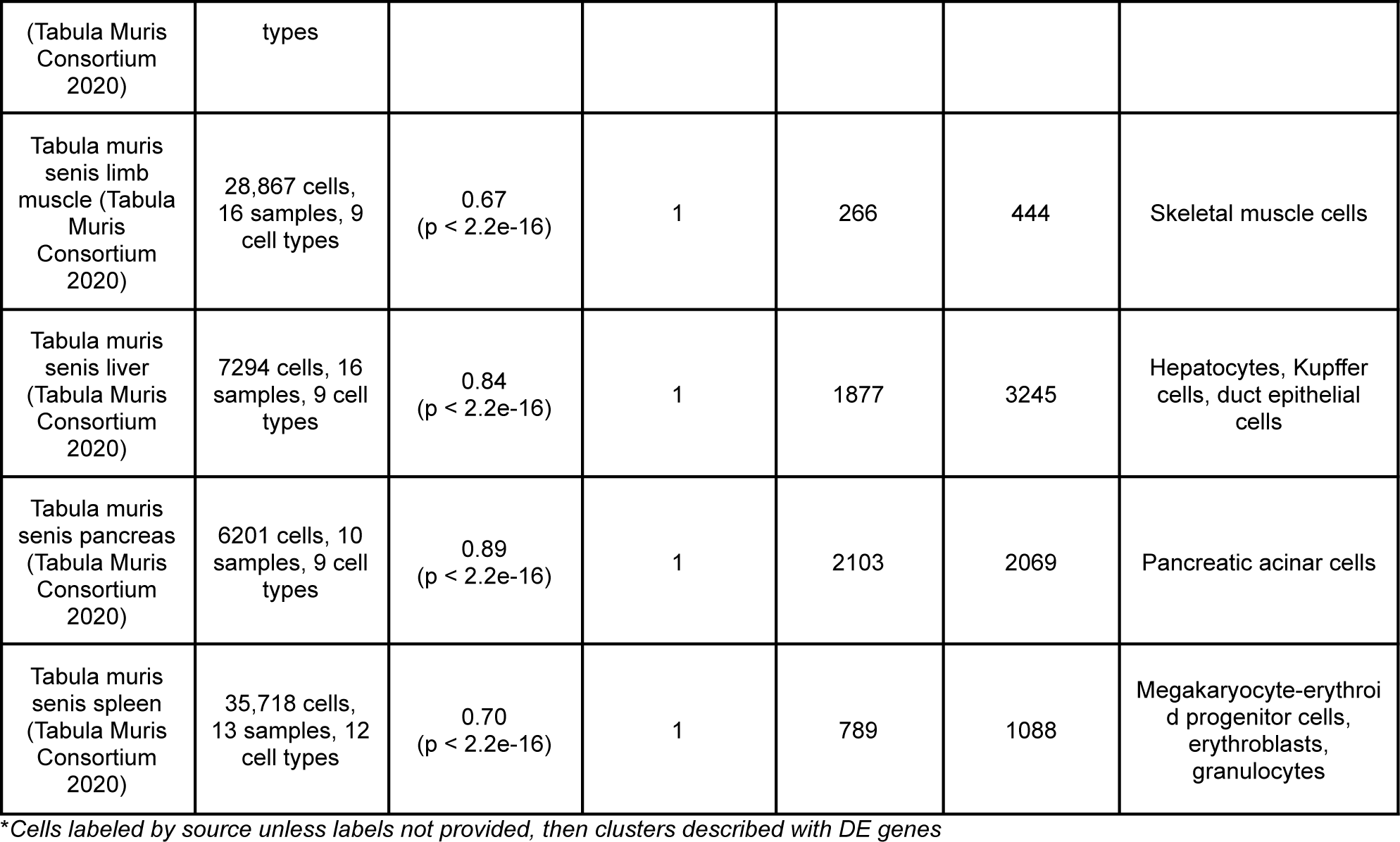
Summary of datasets analyzed with DropletQC.

Other genes that often strongly correlate with the nuclear fraction include those encoding transcripts retained in the nucleus like *NEAT1* and *XIST*. However, these transcripts are not as consistently and strongly expressed as *MALAT1*. As expected, *MALAT1* is often not detected in large proportions of erythrocyte clusters which do not possess a nucleus. Erythrocyte clusters also have a low nuclear fraction.

We also asked if any genes have a strong, negative correlation with the nuclear fraction to indicate high cytoplasmic RNA content. We found no strong patterns, and any negatively correlated genes are often marker genes for poor quality or erythroid cells that did not pass the MALAT1 filter. Examples of these genes include hemoglobin genes, which are markers of erythroid cells lacking a nucleus; hepatocyte-associated genes, as hepatocytes were often found to be poor quality; and mitochondrial genes, which are often highly expressed in stressed or dying cells.

*MALAT1* expression and nuclear fraction values are low in droplets that have other indicators of low quality. This can be seen in the liver, for example, where it is challenging to annotate hepatocyte clusters. Hepatocytes are the most abundant liver cell type, but are also large and proposed to be fragile causing them to become damaged in typical cell isolation protocols (MacParland et al. 2018). As a result, liver single-cell datasets typically suffer from large quantities of ambient RNA expression (Lee et al. 2021), where hepatocyte transcripts are found in every cell type. Clusters that have the highest ambient RNA expression often have low *MALAT1* expression, suggesting that these are likely droplets that have been filled entirely with ambient RNA and may be incorrectly labeled as hepatocytes. In all of the liver datasets we analyzed, hepatocytes were flagged as a problematic cell type based on both nuclear fraction and *MALAT1* expression values (Table 1). Such clusters may also blend into clusters of other cell types, suggesting that droplets may be filled with variable levels of ambient RNA and cell fragments from one or more cell types.

The strong correlation of *MALAT1* with the nuclear fraction, *MALAT1* being a top differentially expressed gene when comparing cells to droplets, and identifying low *MALAT1* values in cells with other indicators of low cell quality suggest that *MALAT1* is a reliable indicator of cell quality in scRNA-seq data.

### Normalized MALAT1 counts can be used to filter out low-quality cells

We next asked if *MALAT1* counts could be used to automatically remove low-quality cells from a scRNA-seq dataset. Analyzing many publicly available datasets, we found that once reads are normalized, *MALAT1* expression typically follows a distribution of expression with at least one major high expression mode with short tails. Therefore, datasets with *MALAT1* expression lower than the short tail of the normal distribution, representing extremely low expression values, can be inferred to be either anucleated cells, empty droplets or droplets containing non-nuclear cell fragments. By analyzing histograms of normalized expression distribution, we can estimate a *MALAT1* expression threshold, below which droplets in the dataset should be flagged for detailed review or removed (Figure 2).

**Figure 2.**
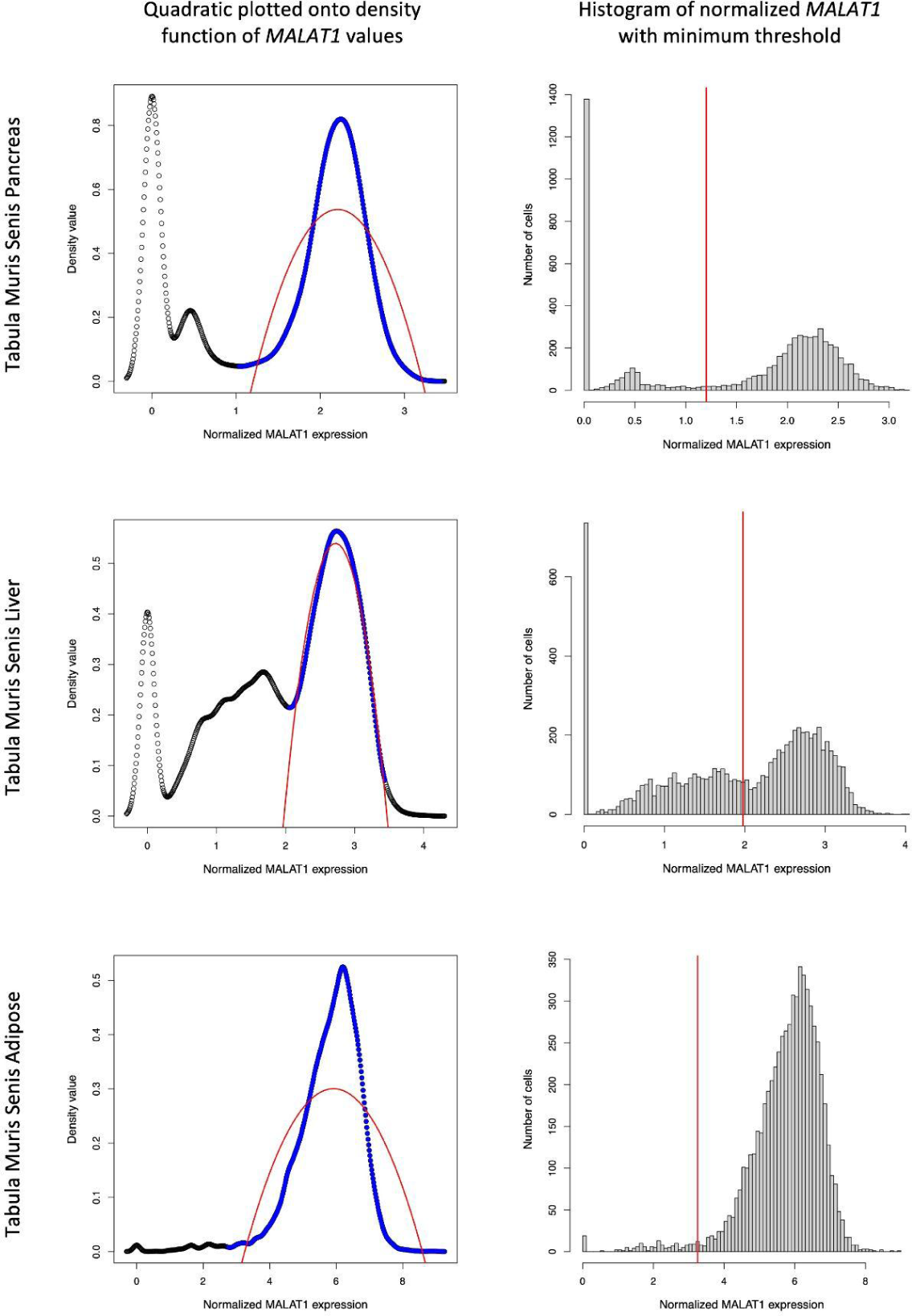
A histogram of normalized *MALAT1* expression can be used to automatically identify a threshold to filter out poor quality cells or empty droplets. The left column represents the density function plotted onto the histogram, with the *MALAT1* peak representing high quality-cells highlighted in blue. The quadratic plotted onto those values is drawn in red, and the lower x-intercept is used to define the *MALAT1* threshold, below which all cells are flagged or removed from the dataset. The right column contains the histogram of normalized *MALAT1* values with the calculated threshold marked as a red vertical line. The y-axis describes the number of cells with the *MALAT1* expression level indicated on the x-axis. The pancreas dataset demonstrates a sample with a high normalized *MALAT1* expression peak, in addition to a low peak below the threshold, and a peak at zero. The liver dataset represents a challenging case, where the sample may have droplets filled with cell fragments as there is no clear divide between the higher *MALAT1* peak and the lower but positive *MALAT1* values. The adipose dataset represents a high-quality sample with very few cells with low *MALAT1* expression.

We created a function to automatically select a threshold for *MALAT1* expression below which droplets are likely to not contain intact cells. The threshold is selected as the lower x-axis intercept of a quadratic function fit over the *MALAT1* expression distribution (Figure 2). We use a quadratic function as a heuristic to easily identify an x-axis intercept that we can use as a rough threshold and avoid modeling the data distribution, which is not generally characterized for scRNA-seq data. Removing cells with lower than this threshold *MALAT1* expression value removes similar cells to DropletQC (Figure 3) and many or all erythroid cells.

**Figure 3.**
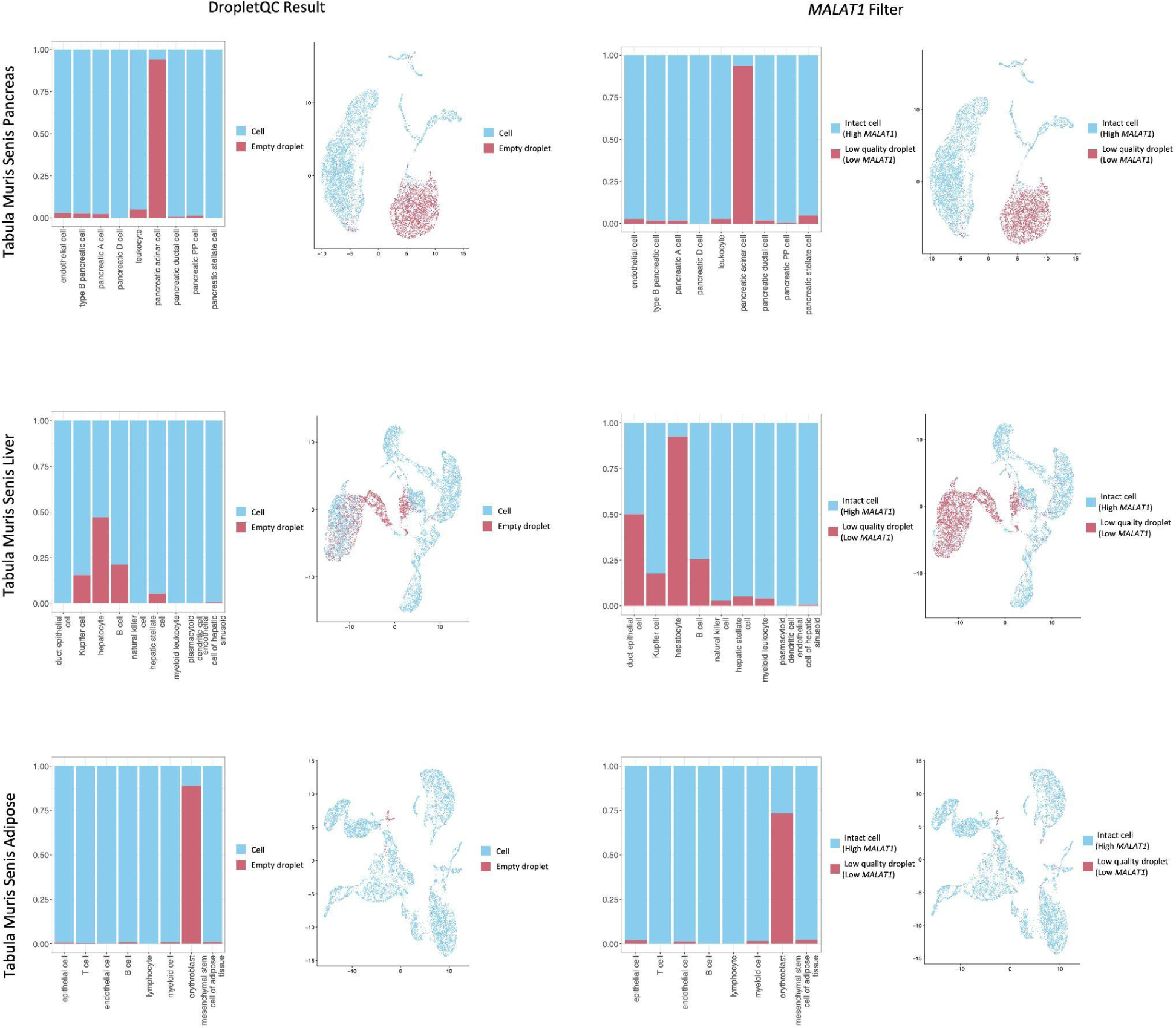
Barplots and UMAPs of cells that have been identified as cells or empty droplets by DropletQC or a *MALAT1* threshold. Cells passing the filters are in blue, while droplets that have not passed the filters are in red. In the barplots, cells are grouped by cell type as identified by the original dataset authors. Precise numbers of filtered cells are found in Table 1. The pancreas dataset represents a typical case, where the DropletQC and *MALAT1* threshold results align closely as there are clear high and low *MALAT1* peaks. The liver sample represents a challenging case, where more moderate *MALAT1* values make it difficult to automatically assign an accurate threshold and expert intervention may be required. The adipose dataset represents a high-quality sample with few low *MALAT1* values, all belonging to erythroblasts.

In some datasets there is no clear distinction between normal and low *MALAT1* values, causing DropletQC and *MALAT1*-based filtering results to differ. This occurs in two integrated liver datasets, where hepatocytes frequently express low to moderate *MALAT1* values with strong batch effects (Table 1, Figure 2). Thus, manual expert intervention is required to verify and potentially adjust our automatically identified *MALAT1* threshold.

### Certain cell types have consistently low or variable MALAT1 expression

We next asked if specific cell types have characteristic *MALAT1* expression distributions. We examined *MALAT1* expression across four different cell atlases from human and mouse: Tabula Sapiens (Tabula Sapiens Consortium et al. 2022) (483,152 cells, 45 tissues), a human cell atlas of fetal gene expression, 1 million cell subset (Cao et al. 2020) (1,001,288 cells, 15 tissues), Tabula Muris Senis 10x (245,389 cells, 16 tissues) (Tabula Muris Consortium 2020), and Tabula Muris Senis Smart-seq2 (110,824 cells, 22 tissues) (Tabula Muris Consortium 2020). We noticed that the same cell types frequently have low *MALAT1* expression, including pancreatic acinar cells, hepatocytes, cardiac muscle cells, and Kupffer cells (Figure 4). Proportions of cells within a cell type with low *MALAT1* expression varies by sample. In some cases, entire populations of cells are removed by both DropletQC and our *MALAT1* threshold filter. In one kidney dataset, 67% of cells may have lost their nuclei (Table 1). Erythroid cells also consistently have low *MALAT1* values, which serves as a positive control for our model, as erythroblasts expel their nuclei during terminal maturation (Menon and Ghaffari 2021). The cells with low *MALAT1* expression that should contain a nucleus (i.e. non-erythroid cells) often represent cells that are challenging to work with because they are large, fragile, or have some other physical characteristic that causes them to become damaged in the single-cell experiment.

**Figure 4.**
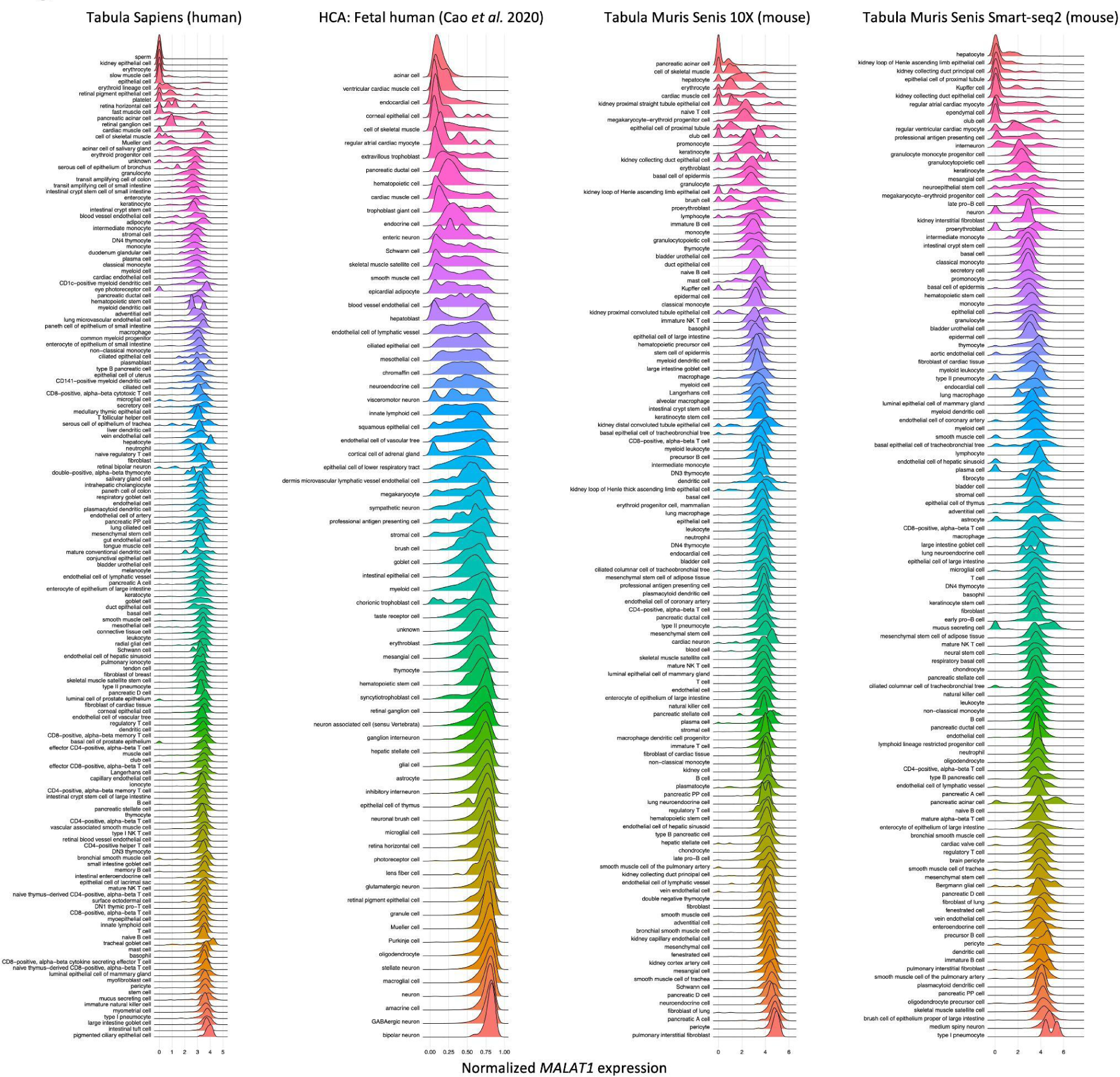
Ridge plots of *MALAT1* expression across cell types identified by four different cell atlases, two human and two mouse: Tabula Sapiens (Tabula Sapiens Consortium et al. 2022), a human cell atlas of fetal gene expression (Cao et al. 2020), Tabula Muris Senis 10x Genomics data (Tabula Muris Consortium 2020), and Tabula Muris Senis Smart-seq2 data (Tabula Muris Consortium 2020). Ridge plots are coloured by cell type. Peak expression values on the left of the plot represent proportions of cells of a given type that may not contain a nucleus, such as an enucleated cell, an empty droplet, or a cell fragment.

In a single dataset, there may be multiple clusters of cells of the same type, but only one of these clusters has low *MALAT1* expression (represented by cell types with two peaks in Figure 4). This suggests that a subset of these cells may have been damaged with their cytoplasmic RNA released into the solution. In this example, the high *MALAT1* cluster may contain normal cells or whole nuclei that have lost some of their cytoplasm, whereas the low *MALAT1* cluster likely represents a group of empty droplets filled with the released cytoplasmic RNA coming from damaged cells that should be removed from the experiment.

## Discussion

Single-cell RNA sequencing maps are being published at a rapid rate. We found that analyzing the *MALAT1* expression distribution across all cells in an experiment is a quick and effective way to identify damaged cells or empty droplets, which helps improve the interpretation of these maps. We developed a fast heuristic that doesn’t rely on splice ratios or BAM files to filter or flag cells with low *MALAT1* expression that can easily be incorporated into a QC filtering pipeline. We tested our method on both human and mouse scRNA-seq data, however our method should be effective across jawed vertebrates and birds where *MALAT1* is conserved (Weghorst et al. 2024).

We compared normalized *MALAT1* expression of both individual samples and integrated datasets to the cells’ nuclear fractions. Although this is effective, for most accurate results, it is best to look at *MALAT1* expression for each sample separately, since technical factors can vary by batch. Normalized *MALAT1* expression is highly ranked when correlated with the nuclear fraction in all datasets we examined except the Tabula Muris Senis bone marrow dataset, where the correlation ranking is 70 (Table 1). In this dataset, there are few cells with low normalized *MALAT1* expression, and almost all are granulocytes. In this dataset, the nuclear fraction is also correlated with any genes not highly expressed in that cluster but strongly expressed in all other cell types (in this case, many ribosomal genes). Although *MALAT1* is also correlated with the nuclear fraction, the ranking is lower since strong correlations exist with multiple genes that happen to not be expressed in granulocytes.

Since technical variation can vary by individual sample and the types of cells within a tissue, each dataset may have its own nuances that guide how it should be treated. For instance, datasets where low *MALAT1* values are largely found in erythroid cells likely do not need additional filtering, as this will simply remove these cells. In addition, it may be difficult to analyze tissues with one predominantly troublesome cell type if this cell type is frequently filtered out of the dataset (e.g. liver hepatocytes). In such cases, it may be better to flag the cells as potentially damaged, and continue to look for broad patterns in the data to interpret. As another example, particularly long cell types like neurons or myocytes, may frequently be captured in fragments that exclude nuclei; however, depending on the importance of capturing nuclear transcripts in these cell types, analyzing a cytoplasmic piece of the cell may be sufficient so long as the interpreter is aware that the nucleus is missing. *MALAT1* expression profiles may also help identify cell types frequently expressing low *MALAT1*, which may need to be treated differently in experimental protocols to prevent them from becoming damaged.

Interestingly, certain datasets contain clusters with high *MALAT1* expression that fall outside of the typical distribution (e.g. the hepatocytes in the Tabula Sapiens atlas in Figure 4). These clusters may represent damaged cells that have lost cytoplasmic RNA, cells with large or multiple nuclei, or cells that may be actively transcribing new genes. We do not use *MALAT1* expression alone to identify cells with whole nuclei but released cytoplasm, as it is challenging to distinguish between the aforementioned scenarios. While methods like DropletQC (Muskovic and Powell 2021) predict cells that have lost cytoplasm based on the nuclear fraction, it is difficult to tell if the abnormal population is the one with high *MALAT1* expression or moderate *MALAT1* expression; an expert may be required to make such a decision. Additional factors support droplet filtering based on the *MALAT1* distribution. For example, when looking at *MALAT1* expression in single-nucleus RNA-seq, *MALAT1* consistently follows a unimodal distribution (Montserrat-Ayuso and Esteve-Codina 2024). Also, when examining raw datasets of all droplets before any filtering, *MALAT1* expression level alone can isolate cells from droplets, yielding results comparable to the current cell isolation bioinformatics pipeline based on a knee plot or removing empty droplets using a background RNA expression distribution. This further supports the idea that *MALAT1* is a reliable measure to discern cells from empty droplets.

Considering many datasets likely contain empty, contaminated droplets and are not yet analyzed for quality using a measure to look for the presence of a nucleus in a droplet, we propose that this easy, quick check should become a standard cell filtering criteria for single-cell data, especially during the iterative process of cell annotation. Further, traditional strict mitochondrial content filters may incorrectly remove healthy cells with high metabolic activity from a dataset (Montserrat-Ayuso and Esteve-Codina 2024); such cells would express moderate to high *MALAT1* levels while having a high mitochondrial content. Cells with high mitochondrial content would still be flagged in a dataset if they were paired with low *MALAT1* expression. This suggests that *MALAT1* expression may serve as an improved QC metric that can potentially replace previous filters, reducing both false positive and false negative rates. Generally, we conclude that *MALAT1* is useful as a technical marker, and should be investigated to aid the quality control of other RNA-based data types to ensure whole nuclei are being captured.

## Methods

For the comparison of DropletQC (v0.0.0.9000) results and normalized *MALAT1* expression levels, we downloaded 16 publicly available datasets, including ten integrated single-cell maps and six single-sample datasets, overall consisting of 118 samples and 215,397 cells across ten healthy tissues, three different disease types, and two species. Datasets were chosen based on the ease of downloading BAM files and the variability of *MALAT1* expression if the dataset was available as an interactive online map.

For 10x Genomics datasets, we downloaded the BAM files and raw and filtered count matrices from https://www.10xgenomics.com/datasets (accessed 2024, May 20). We used EmptyDrops (Lun et al. 2019) on the raw counts matrix and recorded cells versus droplets. Then, we loaded the filtered count matrix into Seurat (v5.0.0), and used their SCTransform pipeline to normalize the data (Hafemeister and Satija 2019), and clustered the datasets at a 0.4 resolution. Differentially expressed (DE) genes were calculated with Presto (v1.0.0) using default parameters to determine the likely cell types for certain clusters (Korsunsky et al. 2019).

For these datasets, we also isolated a subset of droplets from the raw count matrix provided by Cell Ranger and added this to the Seurat object. To do this, we isolated all droplets from the raw count matrix, and isolated cells that contained a minimum number of UMIs that varied depending on the sample (i.e. samples with generally lower UMI counts in the raw count matrix would have a lower minimum UMI threshold). We executed the same normalization and clustering pipeline, and calculated DE genes using Presto with cells divided into three groups: cells classified as a cell by Cell Ranger and DropletQC; cells classified as a cell by Cell Ranger but a droplet by DropletQC; and cells classified as a droplet by Cell Ranger and therefore not included in the original sample.

Samples with available UMAP coordinates were downloaded with the coordinates and normalized count matrices preserved. Therefore no new clustering or normalization was performed, so the values we used match those of the published datasets.

On all datasets, DropletQC was used to calculate the nuclear fraction for each cell when BAM files were available. Cells from samples with missing or improperly formatted BAM files were removed from the dataset. This occurred with two integrated datasets, where some samples had unusable or missing BAM files (unresolved after corresponding with the original authors), leading us to calculate the nuclear fraction for only a subset of cells. DropletQC’s function to classify empty droplets from cells was run on each sample individually using default parameters.

For the *MALAT1* analysis, normalized *MALAT1* values were used and plotted on a histogram. These values were visualized at a dataset rather than sample resolution (i.e. integrated samples were analyzed as a single dataset) as analyzing samples individually was often unnecessary considering the consistency of the pattern, and would require extra steps.

Pearson correlation was used to determine genes correlated with the nuclear fraction and normalized gene expression values. The correlation was calculated across all cells in the dataset that had non-NA nuclear fraction values, which were added when usable BAM files weren’t available to calculate the nuclear fraction.

All figures were created in R (v4.3.1) with packages Seurat (v5.0.0) (Hafemeister and Satija 2019), ggplot2 (v3.4.4) (Wickham 2011), dplyr (v1.1.4) (Wickham et al.), rcartocolor (v2.1.1) (Nowosad), scCustomize (v1.1.3) (Marsh SE (2021) 2024), and viridis (0.6.5) (Garnier et al. 2023); stringr (v1.5.0) (Wickham) was used for formatting object metadata.

## Data and code availability

The *MALAT1* threshold function is available at: https://github.com/BaderLab/MALAT1_threshold

Analysis scripts are at: https://github.com/14zac2/MALAT1Analysis

All data in this study are publicly available at the following links or accession numbers: Human PBMCs from 10x Genomics (10x Genomics 2020d): https://support.10xgenomics.com/single-cell-gene-expression/datasets/4.0.0/Parent_NGSC3_DI_PBMC

Human Hodgkins lymphoma from 10x Genomics (10x Genomics 2020b): https://support.10xgenomics.com/single-cell-gene-expression/datasets/4.0.0/Parent_NGSC3_DI_HodgkinsLymphoma

Human ductal carcinoma from 10x Genomics (10x Genomics 2021): https://www.10xgenomics.com/datasets/7-5-k-sorted-cells-from-human-invasive-ductal-carcinoma-3-v-3-1-3-1-standard-6-0-0

Human glioblastoma from 10x Genomics (10x Genomics 2020c): https://support.10xgenomics.com/single-cell-gene-expression/datasets/4.0.0/Parent_SC3v3_Human_Glioblastoma

Human liver map (MacParland et al. 2018): GSE115469

Human kidney map (Stewart et al. 2019): https://data.humancellatlas.org/explore/projects/abe1a013-af7a-45ed-8c26-f3793c24a1f4 or https://zenodo.org/records/3245841 (Seven available BAM files: CZIKidney7587405, CZIKidney7587406, CZIKidney7587407, CZIKidney7587408, CZIKidney7632802, CZIKidney7632803, CZIKidney7632804).(Seven available BAM files: CZIKidney7587405, CZIKidney7587406, CZIKidney7587407, CZIKidney7587408, CZIKidney7632802, CZIKidney7632803, CZIKidney7632804).

Mouse cortex from 10x Genomics (10x Genomics 2020a): https://support.10xgenomics.com/single-cell-gene-expression/datasets/4.0.0/SC3_v3_NextGem_DI_Neuron_10K?

Mouse heart from 10x Genomics (10x Genomics 2018): https://www.10xgenomics.com/datasets/1-k-heart-cells-from-an-e-18-mouse-v-3-chemistry-3-standard-3-0-0

Human scRNA-seq liver map (Andrews et al. 2022): GSE185477

All Tabula Muris Senis datasets (Tabula Muris Consortium 2020): GSE132042 and https://registry.opendata.aws/tabula-muris-senis/ and https://cellxgene.cziscience.com/collections/0b9d8a04-bb9d-44da-aa27-705bb65b54eb. All BAM files collected except Tabula Muris Senis Adipose. All BAM files collected except Tabula Muris Senis Adipose sample MACA_18m_M_SCAT_53.sample MACA_18m_M_SCAT_53.

Tabula Sapiens (Tabula Sapiens Consortium et al. 2022): https://cellxgene.cziscience.com/collections/e5f58829-1a66-40b5-a624-9046778e74f5 and https://registry.opendata.aws/tabula-sapiens/

A human cell atlas of fetal gene expression (Cao et al. 2020): GSE156793 and https://cellxgene.cziscience.com/collections/c114c20f-1ef4-49a5-9c2e-d965787fb90c

## Notes

https://support.10xgenomics.com/single-cell-gene-expression/datasets/4.0.0/Parent_NGSC3_DI_PBMC

https://support.10xgenomics.com/single-cell-gene-expression/datasets/4.0.0/Parent_NGSC3_DI_HodgkinsLymphoma

https://www.10xgenomics.com/datasets/7-5-k-sorted-cells-from-human-invasive-ductal-carcinoma-3-v-3-1-3-1-standard-6-0-0

https://support.10xgenomics.com/single-cell-gene-expression/datasets/4.0.0/Parent_SC3v3_Human_Glioblastoma

https://www.ncbi.nlm.nih.gov/geo/query/acc.cgi?acc=GSE115469

https://data.humancellatlas.org/explore/projects/abe1a013-af7a-45ed-8c26-f3793c24a1f4

https://support.10xgenomics.com/single-cell-gene-expression/datasets/4.0.0/SC3_v3_NextGem_DI_Neuron_10K?

https://www.10xgenomics.com/datasets/1-k-heart-cells-from-an-e-18-mouse-v-3-chemistry-3-standard-3-0-0

https://registry.opendata.aws/tabula-muris-senis/

https://cellxgene.cziscience.com/collections/e5f58829-1a66-40b5-a624-9046778e74f5

### Competing Interest Statement

The authors have declared no competing interest.

### Summary of Updates

We have cited additional manuscripts and commented on their work.

